# pulse2percept: A Python-based simulation framework for bionic vision

**DOI:** 10.1101/148015

**Authors:** Michael Beyeler, Geoffrey M. Boynton, Ione Fine, Ariel Rokem

## Abstract

By 2020 roughly 200 million people worldwide will suffer from photoreceptor diseases such as retinitis pigmentosa and age-related macular degeneration, and a variety of retinal sight restoration technologies are being developed to target these diseases. One technology, analogous to cochlear implants, uses a grid of electrodes to stimulate remaining retinal cells. Two brands of retinal prostheses are currently approved for implantation in patients with late stage photoreceptor disease. Clinical experience with these implants has made it apparent that the vision restored by these devices differs substantially from normal sight. To better understand the outcomes of this technology, we developed *pulse2percept*, an open-source Python implementation of a computational model that predicts the perceptual experience of retinal prosthesis patients across a wide range of implant configurations. A modular and extensible user interface exposes the different building blocks of the software, making it easy for users to simulate novel implants, stimuli, and retinal models. We hope that this library will contribute substantially to the field of medicine by providing a tool to accelerate the development of visual prostheses.

## 1 Introduction

Two frequent causes of blindness in the developed world are age-related macular degeneration (AMD) and retinitis pigmentosa (RP) [BBB+84], [Gro04]. Both of these diseases have a hereditary component, and are characterized by a progressive degeneration of photoreceptors in the retina that lead to gradual loss of vision.

Microelectronic retinal prostheses have been developed in an effort to restore sight to RP and AMD patients. Analogous to cochlear implants, these devices function by electrically stimulating surviving retinal neurons in order to evoke neuronal responses that are transmitted to the brain and interpreted by patients as visual percepts (Fig. 1). Two of these devices are already approved for commercial use, and a number of other companies have either started or are planning to start clinical trials of devices in the near future. Other types of technologies, such as optogenetics and genetic modification are also areas of active research. Blinded individuals may potentially be offered a wide range of sight restoration options within a decade [FCL15].

**Fig. 1:**
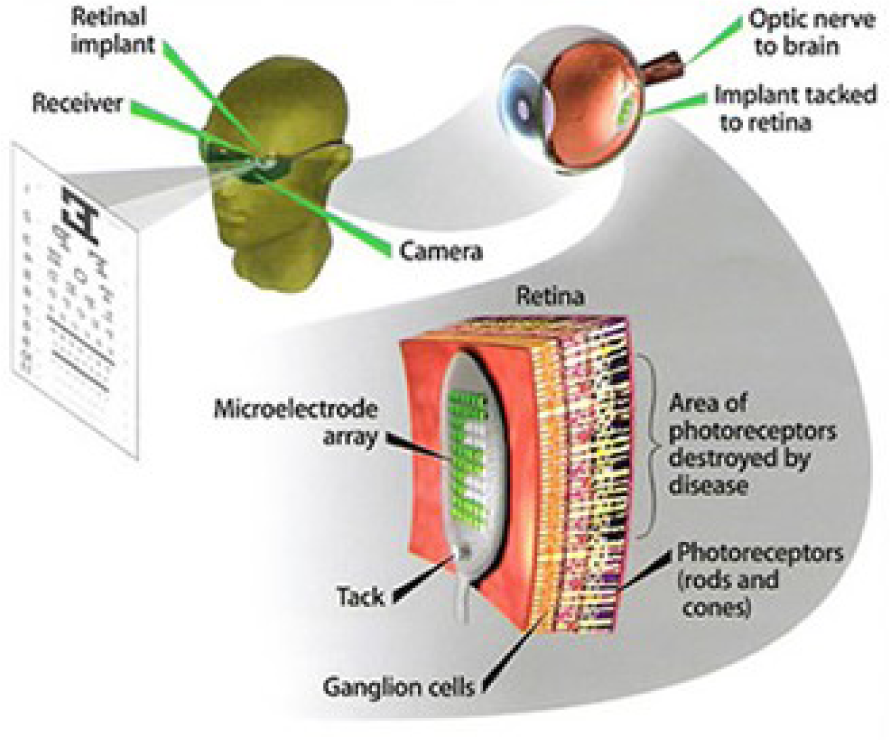
Electronic retinal prosthesis. Light from the visual scene is captured by an external camera and transformed into electrical pulses delivered through microelectrodes to stimulate the retina.

One major challenge in the development of retinal prostheses is predicting what patients will see when they use their devices. Interactions between implant electronics and the underlying neurophysiology cause nontrivial perceptual distortions in both space and time [FB15], [BRBFss] that severely limit the quality of the generated visual experience.

Our goal was to develop a simulation framework that could describe the visual percepts of retinal prosthesis patients over space and time. We refer to these simulations as ‘virtual patients’, analogous to the virtual prototyping that has proved so useful in other complex engineering applications.

Here we present an open-source implementation of these models as part of *pulse2percept*, a BSD-licensed Python-based simulation framework [BSD17] that relies on the NumPy and SciPy stacks, as well as contributions from the broader Python community. Based on the detailed specification of a patient’s implant configuration, and given a desired electrical stimulus, the model predicts the perceptual distortions experienced by ‘virtual patients’ over both space and time.

We hope that this library will contribute substantially to the field of medicine by providing a tool to accelerate the development of visual prostheses. Researchers may use this tool to improve stimulation protocols of existing implants or to aid development of future devices. In addition, this tool might guide government agencies, such as the FDA and Medicare, in making reimbursement decisions. Furthermore, this tool can be used to guide patients and doctors in their decision as to when or whether to be implanted, and which device to select.

The remainder of this paper is organized as follows: We start by introducing the neuroscience background necessary to understand the interactions between implant electronics and the underlying neurophysiology. We then detail the computational model that underlies *pulse2percept*, before we give a simple usage example and go into implementation details. We then review our solutions to various technical challenges, and conclude by discussing the broader impact for this work for computational neuroscience and neural engineering communities in more detail.

## Background

The first steps in seeing begin in the retina, where a mosaic of photoreceptors converts incoming light into electrochemical signals that encode the intensity of light as a function of position (two dimensions), wavelength, and time [Rod98]. The electrochemical signal is passed on to specialized neuronal circuits consisting of a variety of cell types (such as bipolar, amacrine, and horizontal cells), which act as feature detectors for basic sensory properties, such as spatial contrast and temporal frequency. These sensory features are then encoded in parallel across approximately 1.5 million retinal ganglion cells, which form the output layer of the retina. Each ganglion cell relays the electrical signal to the brain via a long axon fiber that passes from the ganglion cell body to the optic nerve and on to the brain.

Diseases such as RP and AMD are characterized by a progressive degeneration of photoreceptors, gradually affecting other layers of the retina [HPdJ^+^99], [MNS08]. In severe end-stage RP, roughly 95% of photoreceptors, 20% of bipolar cells, and 70% of ganglion cells degenerate [SHd^+^97]. In addition, the remaining cells undergo corruptive re-modeling in late stages of the disease [MJWS03], [MJ03], so that little or no useful vision is retained.

Microelectronic retinal prostheses have been developed in an effort to restore sight to individuals suffering from RP or AMD. Analogous to cochlear implants, the goal of retinal prostheses is to electrically stimulate surviving bipolar or ganglion cells in order to evoke neuronal responses that are interpreted by the brain as visual percepts. The electrical stimulus delivers charge to the cell membrane that depolarizes the neuron and opens voltage-sensitive ion channels. This bypasses the natural presynaptic neurotransmitter excitation and causes the activated neurons to stimulate their postsynaptic targets. Therefore, selective spatiotemporal modulation of retinal neurons with an array of electrodes should allow a prosthesis to coordinate retinal activity in place of natural photoreceptor input [WWH16].

Several types of retinal prostheses are currently in development. These vary in their user interface, light-detection method, signal processing, and microelectrode placement within the retina (for a recent review see [WWH16]). As far as our model is concerned, the critical factor is the placement of the microelectrodes on the retina (Fig. 2). The three main locations for microelectrode implant placement are *epiretinal* (i.e., on top of the retinal surface, above the ganglion cells), *subretinal* (i.e., next to the bipolar cells in the space of the missing photoreceptors), and *suprachoroidal* (i.e., between the choroid and the sclera). Each of these approaches is similar in that light from the visual scene is captured via a camera and transformed into electrical pulses delivered through electrodes to stimulate the retina.

**Fig. 2:**
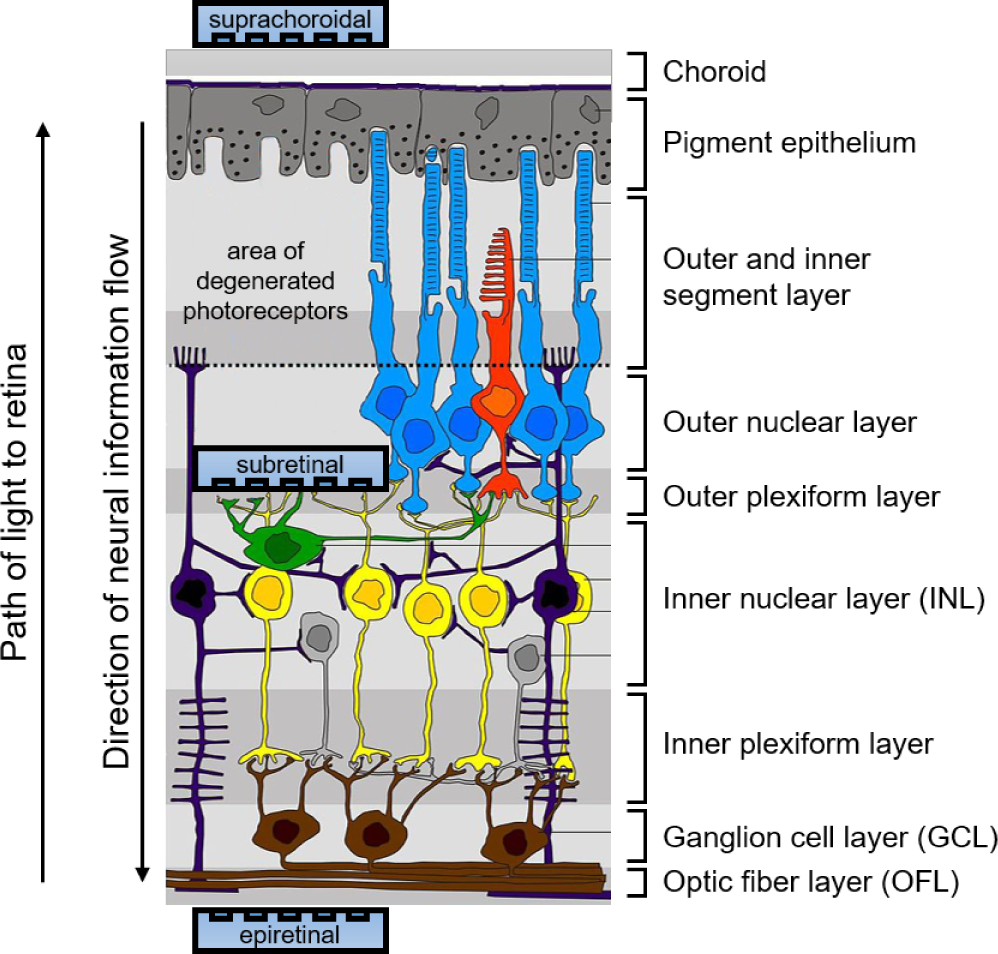
Diagram of the retina and common locations of retinal prosthesis microelectrode arrays. Retinitis pigmentosa causes severe photoreceptor degeneration. Epiretinal electrode arrays are placed in the vitreous, next to the optic fiber layer (OFL). Subretinal arrays are placed by creating a space between the choroid and remaining retinal tissue. Suprachoroidal arrays are placed behind the choroid. pulse2percept allows for simulation of processing in the inner nuclear layer (INL), ganglion cell layer (GCL), and optic fiber layer (OFL). Based on “Retina layers” by Peter Hartmann, CC BY-SA 3.0.

As mentioned above, two devices are currently approved for commercial use and are being implanted in patients across the US and Europe: the Argus II device (epiretinal, Second Sight Medical Products Inc., [dCDH^+^16]) and the Alpha-IMS system (subretinal, Retina Implant AG, [SBSB^+^15]). At the same time, a number of other companies have either started or are planning to start clinical trials in the near future, potentially offering a wide range of sight restoration options for the blind within a decade [FCL15].

However, clinical experience with existing retinal prostheses makes it apparent that the vision provided by these devices differs substantially from normal sight. Interactions between implant electronics and the underlying neurophysiology cause nontrivial perceptual distortions in both space and time [FB15], [BRBFss] that severely limit the quality of the generated visual experience. For example, stimulating a single electrode rarely produces the experience of a ‘dot’ of light, instead leading to percepts that vary drastically in shape, varying in description from ‘blobs’, to ‘streaks’ and ‘half-moons’. The percept produced by stimulating a single electrode with a continuous pulse train also fades over time, usually disappearing over a course of seconds. As a result, patients do not report seeing an interpretable world. One patient describes it as like: *“… looking at the night sky where you have millions of twinkly lights that almost look like chaos”* [Pre15].

Previous work by our group has focused on development of computational models to describe some of these distortions for a small number of behavioral observations in either space [NFH^+^12] or time [HGW^+^09]. Here we present a model that can describe spatial distortions, temporal nonlinearities, and spatiotemporal interactions reported across a wide range of conditions, devices, and patients.

**Fig. 3:**
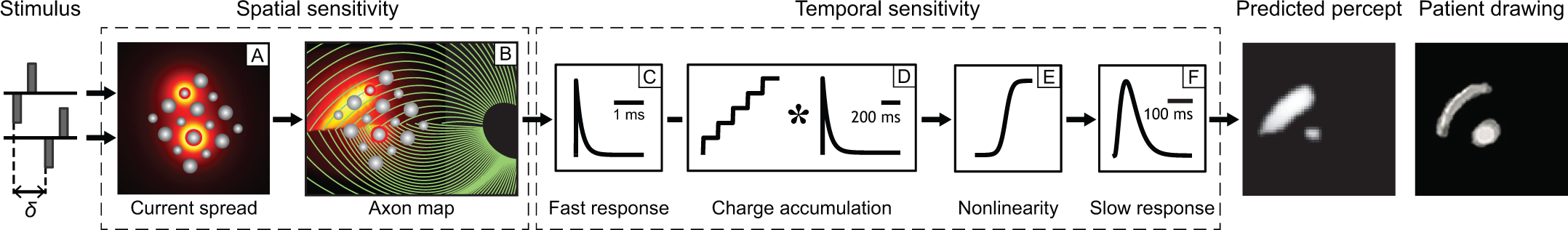
Full model cascade. A simulated electrical stimulus is processed by a series of linear filtering and nonlinear processing steps that model the spatial (A, B) and temporal sensitivity (C-F) of the retinal tissue. An Argus I device is shown (16 electrodes of 260 or 520 microns diameter arranged in a checkerboard pattern). The output of the model is a prediction of the visual percept in both space and time (example frame shown), which can be compared to human patients’ drawings.

## Computational Model of Bionic Vision

Analogous to models of cochlear implants [Sha89], the goal of our computational model is to approximate, via a number of linear filtering and nonlinear processing steps, the neural computations that convert electrical pulse trains across multiple electrodes into a perceptual experience in space and time.

Model parameters were chosen to fit data from a variety of behavioral experiments in patients with prosthetic devices. For example, in threshold experiments patients were asked to report whether or not they detected a percept. Across many trials, the minimum stimulation current amplitude needed to reliably detect the presence of a percept on 80% of trials was found. This threshold was measured across a variety of pulse trains that varied across dimensions such as frequency, duty cycle, and duration. In other experiments patients reported the apparent brightness or size of percepts on a rating scale. In others patients drew the shapes of the percepts evoked by stimulation. The model has been shown to generalize across individual electrodes, patients, and devices, as well as across different experiments. Detailed methods of how the model was validated can be found in [HGW^+^09], [NFH^+^12], [BRBFss]. Here we provide a brief overview of the model cascade.

The full model cascade for an Argus I epiretinal prosthesis is illustrated in Fig. 3. The Argus I device simulated here consists of electrodes of 260*µm* and 520*µm* diameter, arranged in a checkerboard pattern (Fig. 3 A). In this example, input to the model is a pair of simulated pulse trains phase-shifted by *δ* ms, which are delivered to two individual simulated electrodes.

The first stages of the model describe the spatial distortions resulting from interactions between the electronics and the neuroanatomy of the retina. We assume that the current density caused by electrical stimulation decreases as a function of distance from the edge of the electrode [ABK^+^08]:

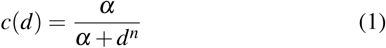

where *d* is the 3D Euclidean distance to the electrode edge, *α* = 14000 and the exponent *n* = 1.69. Current fields for two stimulated electrodes are shown, Fig. 3 A (the hotter the color, the higher the current).

Due to the fact that epiretinal implants sit on top of the optic fiber layer (Fig. 2), they do not only stimulate ganglion cell bodies but also ganglion cell axons. It is critical to note that, perceptually, activating an axon fiber that passes under a stimulated electrode is indistinguishable from the percept that would be elicited by activating the corresponding ganglion cell *body*. The produced visual percept will appear in the spatial location in visual space for which the corresponding ganglion cell and axon usually encodes information. Ganglion cells send their axons on highly stereotyped pathways to the optic disc (green lines in Fig. 3 B), which have been mathematically described [JNS^+^09]. As a result, electrical stimulation of axon fibers leads to predictable visual ‘streaks’ or ‘comet trails’ that are aligned with the axonal pathways.

We therefore convert the spatial map of current densities into a tissue activation map by accounting for axonal stimulation. We model the sensitivity of axon fibers as decreasing exponentially as a function of distance from the corresponding ganglion cell bodies. The resulting tissue activation map across the retinal surface is shown as a heatmap in Fig. 3 B (the hotter the color, the larger the amount of tissue stimulation).

The remaining stages of the model describe temporal nonlinearities. Every pixel of the tissue activation map is modulated over time by the applied electrical pulse train in order to predict a perceived brightness value that varies over time. This involves applying a series of linear filtering (Fig. 3 C, D, and F) and nonlinear processing (Fig. 3 E) steps in the time domain that are designed to approximate the processing of visual information within the retina and visual cortex.

Linear responses in Fig. 3 C, D, and F are modeled as temporal low-pass filters, or ‘leaky integrators’, using gamma functions of order *n*:

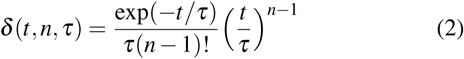

where *t* is time, *n* is the number of identical, cascading stages, and *τ* is the time constant of the filter.

The first temporal processing step convolves the timeseries of tissue activation strengths *f* (*t*) at a particular spatial location with a one-stage gamma function (*n* = 1, time constant *τ*_1_ = 0.42 ms) to model the impulse response function of retinal ganglion cells (Fig. 3 C):

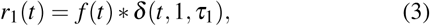

where * denotes convolution.

Behavioral data suggests that the system becomes less sensitive as a function of accumulated charge. This is implemented by calculating the amount of accumulating charge at each point of time in the stimulus, *c*(*t*), and by convolving this accumulation with a second one-stage gamma function (*n* = 1, time constant *τ*_2_ = 45.3 ms; Fig. 3 D). The output of this convolution is scaled by a factor *ε*_1_ = 8.3 and subtracted from *r*_1_ (Eq. 3):

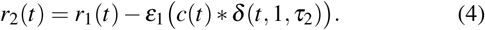

The response *r*_2_(*t*) is then passed through a stationary nonlinearity (Fig. 3 E) to model the nonlinear input-output relationship of ganglion cell spike generation:

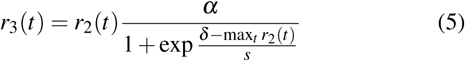

where *α* = 14 (asymptote), *s* = 3 (slope), and *δ* = 16 (shift) are chosen to match the observed psychophysical data.

Finally, the response *r*_3_(*t*) is convolved with another low-pass filter described as a three-stage gamma function (*n* = 3, *τ*_3_ = 26.3 ms) intended to model slower perceptual processes in the brain (Fig. 3 F):

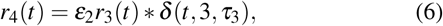

where *ε*_2_ = 1000 is a scaling factor used to scale the output to subjective brightness values in the range [0, 100].

The output of the model is thus a movie of brightness values that corresponds to the predicted perceptual experience of the patient. An example percept generated is shown on the right-hand side of Fig. 3 (‘predicted percept’, brightest frame in the movie).

## Implementation and Results

### Code Organization

The *pulse2percept* project seeks a trade-off between optimizing for computational performance and ease of use. To facilitate ease of use, we organized the software as a standard Python package, consisting of the following primary modules:

- `api`: a top-level Application Programming Interface.
- `implants`: implementations of the details of different retinal prosthetic implants. This includes Second Sight’s Argus I and Argus II implants, but can easily be extended to feature custom implants (see Section Extensibility).
- `retina`: implementation of a model of the retinal distribution of nerve fibers, based on [JNS^+^09], and an implementation of the temporal cascade of events described in Eqs. 2-6. Again, this can easily be extended to custom temporal models (see Section Extensibility).
- `stimuli`: implementations of commonly used electrical stimulation protocols, including methods for translating images and movies into simulated electrical pulse trains. Again, this can easily be extended to custom stimulation protocols (see Section Extensibility).
- `files`: a means to load/store data as images and videos.
- `utils`: various utility and helper functions.

### Basic Usage

Here we give a minimal usage example to produce the percept shown on the right-hand side of Fig. 3.

Convention is to import the main `pulse2percept` module as `p2p`. Throughout this paper, if a class is referred to with the prefix `p2p`, it means this class belongs to the main pulse2percept library (e.g., `p2p.retina`):

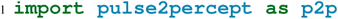

`p2p.implants`: Our goal was to create electrode implant objects that could be configured in a highly flexible manner. As far as placement is concerned, an implant can be placed at a particular location on the retina (`x_center, y_center`) with respect to the fovea (in microns), and rotated as you see fit (rot):

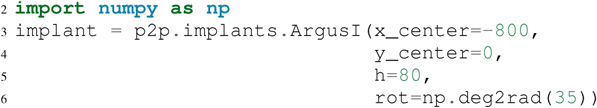

Here, we make use of the `ArgusI` array type, which provides predefined values for array type (‘epiretinal’) and electrode diameters. In addition, the distance between the array and the retinal tissue can be specified via the height parameter (h), either on a per-electrode basis (as a list) or using the same value for all electrodes (as a scalar).

The electrodes within the implant can also be specified. An implant is a wrapper around a list of `p2p.implants.Electrode` objects, which are accessible via indexing or iteration (e.g., via [`for i in implant`]). This allows for specification of electrode diameter, position, and distance to the retinal tissue on a per-electrode basis. Once configured, every Electrode object in the implant can also be assigned a name (in the Argus I implant, they are A1 - A16; corresponding to the names that are commonly used by Second Sight Medical Products Inc.). The first electrode in the implant can be accessed both via its index (`implant[0]`) and its name (`implant[’A1’]`).

Once the implant is created, it can be passed to the simulation framework. This is also where you specify the back end (currently supported are ‘serial’, ‘joblib’ [Job16], and ‘dask’ [Das16]):

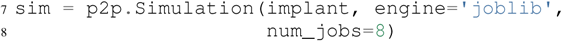

The simulation framework provides a number of setter functions for the different layers of the retina. These allow for flexible specification of optional settings, while abstracting the underlying functionality.

`p2p.retina`: This includes the implementation of a model of the retinal distribution of nerve fibers, based on [JNS^+^09], and implementations of the temporal cascade of events described in Eqs. 2-6.

Things that can be set include the spatial sampling rate of the optic fiber layer (`ssample`) as well as the spatial extent of the retinal patch to be simulated (given by the corner points [`xlo, ylo`] and [`xhi, yhi`]). If the coordinates of the latter are not given, a patch large enough to fit the specified electrode array will be automatically selected:

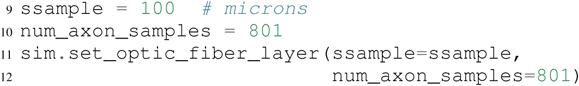

Similarly, for the ganglion cell layer we can choose one of the pre-existing cascade models and specify a temporal sampling rate:

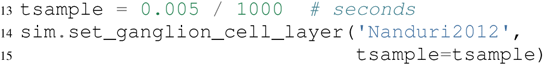

As its name suggest, `‘Nanduri2012’` implements the model detailed in [NFH^+^12]. Other options include `‘Horsager2009’` [HGW^+^09] and ‘latest’.

It’s also possible to specify your own (custom) model, see Section Extensibility below.

`p2p.stimuli`: A stimulation protocol can be specified by assigning stimuli from the p2p.stimuli module to specific electrodes. An example is to set up a pulse train of particular stimulation frequency, current amplitude and duration. Because of safety considerations, all real-world stimuli must be balanced biphasic pulse trains (i.e., they must have a positive and negative phase of equal area, so that the net current delivered to the tissue sums to zero).

It is possible to specify a pulse train for each electrode in the implant as follows:

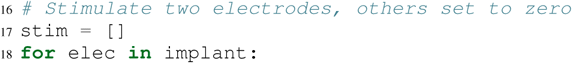

**Fig. 4:**
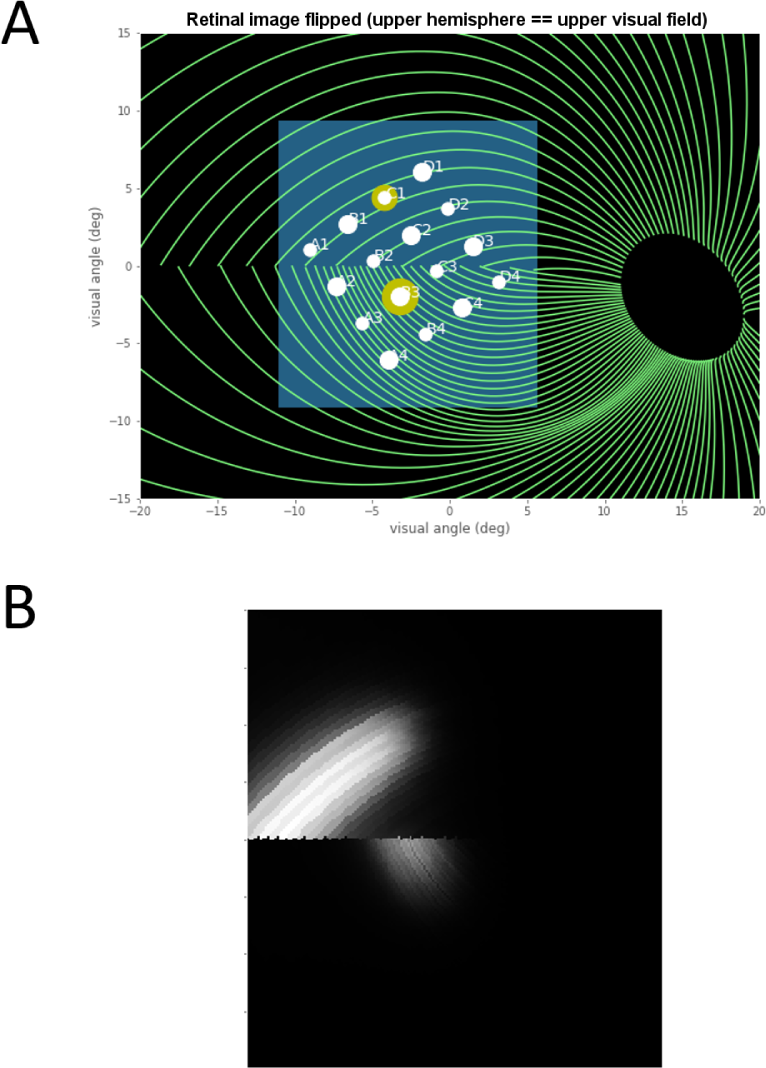
Model input/output generated by the example code. (A) An epiretinal Argus I array is placed near the fovea, and two electrodes (‘C1’ and ‘B3’) are stimulated with 50 Hz, 20 uA pulse trains. The plot is created by lines 34-36. Note that the retinal image is flipped, so that the upper hemisphere corresponds to the upper visual field. (B) Predicted visual percept (example frame shown). The plot is created by line 41.

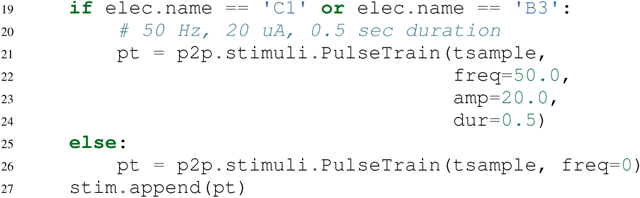

However, since implants are likely to have electrodes numbering in the hundreds or thousands, this method will rapidly become cumbersome when assigning pulse trains across multiple electrodes. Therefore, an alternative is to assign pulse trains to electrodes via a dictionary:

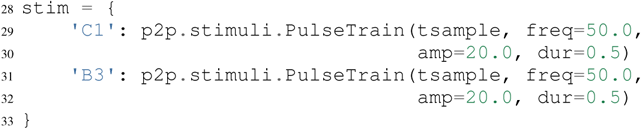

At this point, we can highlight the stimulated electrodes in the array:

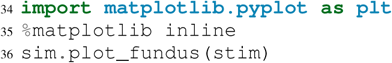

The output can be seen in Fig. 4 A.

Finally, the created stimulus serves as input to sim.pulse2percept, which is used to convert the pulse trains into a percept. This allows users to simulate the predicted percepts for simple input stimuli, such as stimulating a pair of electrodes, or more complex stimuli, such as stimulating a grid of electrodes in the shape of the letter A.

At this stage in the model it is possible to consider which retinal layers are included in the temporal model, by selecting from the following list (see Fig. 2 for a schematic of the anatomy):

- `‘OFL’`: optic fiber layer (OFL), where ganglion cell axons reside,
- `‘GCL’`: ganglion cell layer (GCL), where ganglion cell bodies reside, and
- `‘INL’`: inner nuclear layer (INL), where bipolar cells reside.

A list of retinal layers to be included in the simulation is then passed to the pulse2percept method:

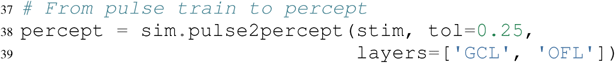

This allows the user to run simulations that include only the layers relevant to a particular simulation. For example, axonal stimulation and the resulting axon streaks can be ignored by omitting `‘OFL’` from the list. By default, all three supported layers are included.

Here, the output `percept` is a `p2p.utils.TimeSeries` object that contains the time series data in its data container. This time series consists of brightness values (arbitrary units in [0, 100]) for every pixel in the percept image.

Alternatively, it is possible to retrieve the brightest (mean over all pixels) frame of the time series:

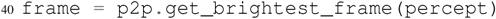

Then we can plot it with the help of Matplotlib (Fig. 4 B):

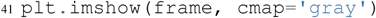

`p2p.files`: *pulse2percept* offers a collection of functions to convert the `p2p.utils.TimeSeries` output into a movie file via scikit-video [Sci17] and ffmpeg [FFm10].

For example, a percept can be stored to an MP4 file as follows:

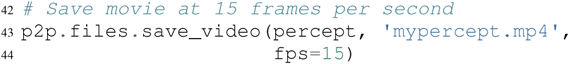

For convenience, *pulse2percept* provides a function to load a video file and convert it to the p2p.utils.TimeSeries format:

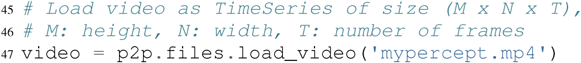

Analogously, *pulse2percept* also provides functionality to go directly from images or videos to electrical stimulation on an electrode array:

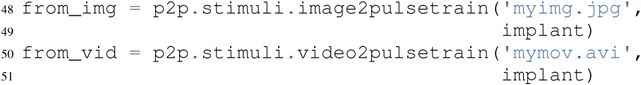

These functions are based on functionality provided by scikit-image [S v14] and scikit-video [Sci17], respectively, and come with a number of options to specify whether brightness should be encoded as pulse train amplitude or frequency, at what frame rate to sample the movie, whether to maximize or invert the contrast, and so on.

### Extensibility

As described above, our software is designed to allow for implants, retinal models, and pulse trains to be customized. We provide extensibility mainly through mechanisms of class inheritance.

Retinal Implants: Creating a new implant involves inheriting from `p2p.implants.ElectrodeArray` and providing a property `etype`, which is the electrode type (e.g., `‘epiretinal’`, `‘subretinal’`):

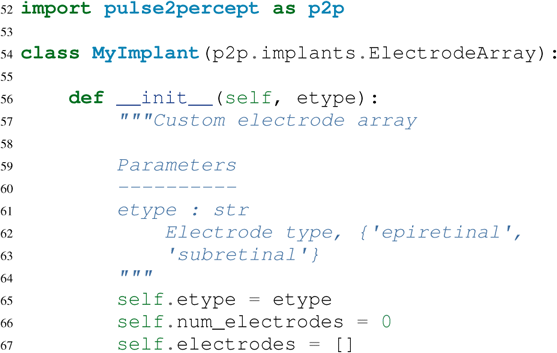

Then new electrodes can be added by utilizing the add_electrodes method of the base class:

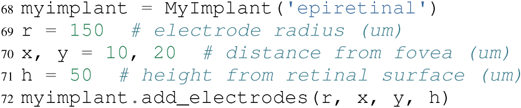

Retinal cell models: Any new ganglion cell model is described as a series of temporal operations that are carried out on a single pixel of the image. It must provide a property called `tsample`, which is the temporal sampling rate, and a method called `model_cascade`, which translates a single-pixel pulse train into a single-pixel percept over time:

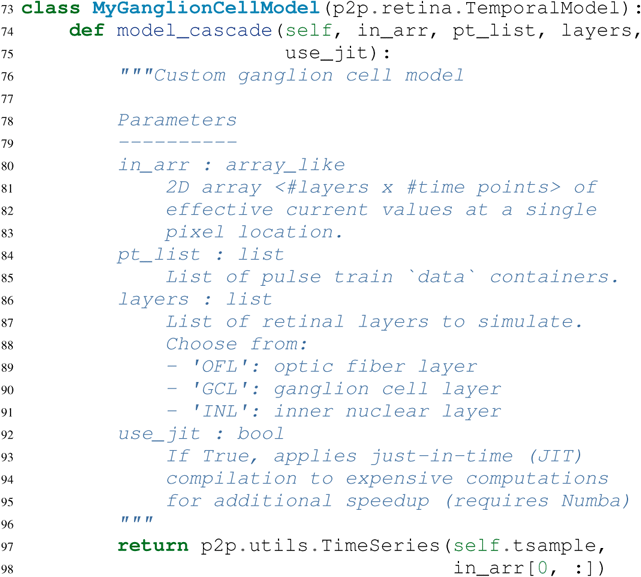

This method can then be passed to the simulation framework:

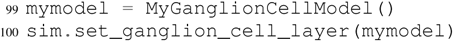

It will then automatically be selected as the implemented ganglion cell model when `sim.pulse2percept` is called.

Stimuli: The smallest stimulus building block provided by *pulse2percept* consists of a single pulse of either positive current (anodic) or negative current (cathodic), which can be created via `p2p.stimuli.MonophasicPulse`. However, as described above, any real-world stimulus must consist of biphasic pulses with zero net current. A single biphasic pulse can be created via p2p.stimuli.BiphasicPulse. A train of such pulses can be created via p2p.stimuli.PulseTrain. This setup gives the user the opportunity to build their own stimuli by creating pulse trains of varying amplitude, frequency, and inter-pulse intervals.

In order to define new pulse shapes and custom stimuli, the user can either inherit from any of these stimuli classes or directly from p2p.utils.TimeSeries. For example, a biphasic pulse can be built from two monophasic pulses as follows:

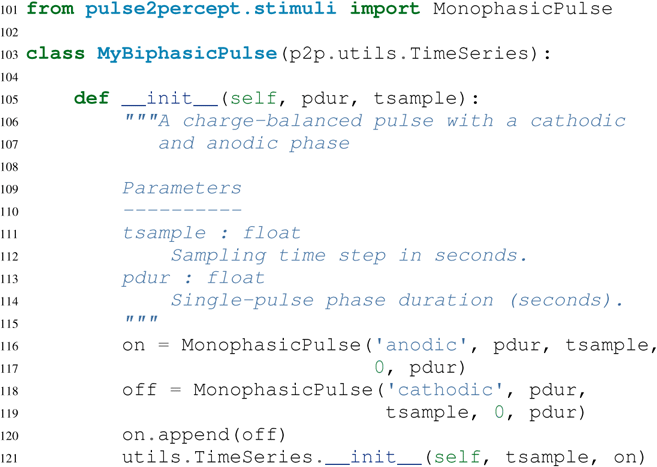

### Implementation Details

*pulse2percept*’s main technical challenge is computational cost: the simulations require a fine subsampling of space, and span several orders of magnitude in time. In the space domain the software needs to be able to simulate electrical activation of individual retinal ganglion cells on the order of microns. In the time domain the model needs to be capable of dealing with pulse trains containing individual pulses on the sub-millisecond time scale that last over several seconds.

Like the brain, we solve this problem through parallelization. Spatial interactions are confined to an initial stage of processing (Fig. 3 A, B), after which all spatial computations are parallelized using two back ends (Joblib [Job16] and Dask [Das16]), with both multithreading and multiprocessing options.

However, even after parallelization, computing the temporal response remains a computational bottleneck. Initial stages of the temporal model require convolutions of arrays (e.g., Eqs. 2 and 3) that describe responses of the model at high temporal resolution (sampling rates on the order of 10 microseconds) for pulse trains lasting for at least 500 milliseconds. These numerically-heavy sections of the code are sped up using a conjunction of three strategies. First, as described above, any given electrode generally only stimulates a subregion of the retina. As a consequence, when only a few electrodes are active, we can often obtain substantial speed savings by ignoring pixels which are not significantly stimulated by any electrode (see tolerance parameter `tol` on line 38 of the example code). Second, electrical stimulation is often carried out at relatively low pulse train frequencies of less than 50 Hz. Since the individual pulses within the pulse train are usually very short (∼75-450 microseconds), input pulse trains tend to be sparse. We can exploit this fact to speed up computation time by avoiding direct convolution with the entire time-series whenever possible, and instead relying on a custom-built sparse convolution function. Preprocessing of sparse pulse train arrays allows us to carry out temporal convolution only for those parts of the time-series that include nonzero current amplitudes. Finally, these convolutions are sped up wih LLVM-base compilation implemented using Numba [LPS15].

**Fig. 5:**
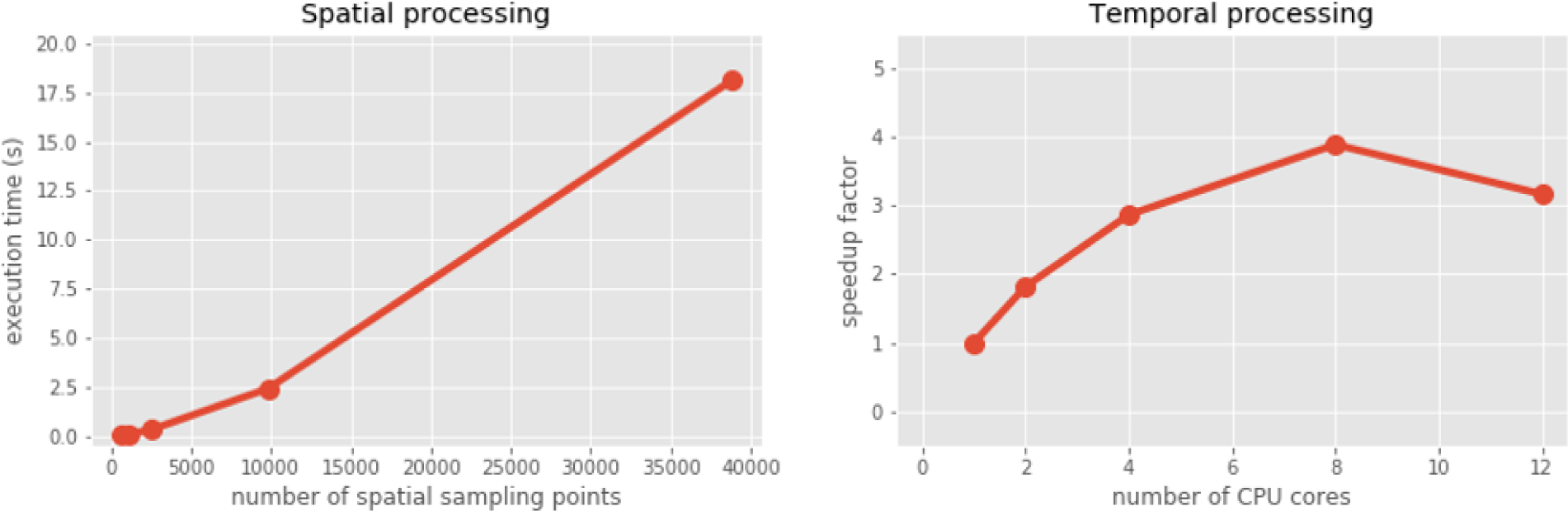
Computational performance. (A) Compute time to generate an ‘effective stimulation map’ is shown as a function of the number of spatial sampling points used to characterize the retina. (B) Speedup factor (serial execution time / parallel execution time) is shown as a function of the number of CPU cores. Execution times were collected for the an Argus II array (60 electrodes) simulating the letter ‘A’ (roughly 40 active electrodes, 20 Hz/20 uA pulse trains) over a period of 500 ms (tsample was 10 microseconds, ssample was 50 microns). Joblib and Dask parallelization back ends gave similar results.

### Computational Performance

We measured computational performance of the model for both spatial and temporal processing using a 12-core Intel Core i7-5930K operating at 3.50 GHz (64GB of RAM).

The initial stages of the model scale as a function of the number of spatial sampling points in the retina, as shown in Fig. 5 A. Since the spatial arrangement of axonal pathways does not depend on the stimulation protocol or retinal implant, *pulse2percept* automatically stores and re-uses the generated spatial map depending on the specified set of spatial parameters.

The remainder of the model is carried out in parallel, with the resulting speedup factor shown in Fig. 5 B. Here, the speedup factor is calculated as the ratio of single-core execution time and parallel execution time. On this particular machine, the maximum speedup factor is obtained with eight cores, above which the simulation shifts from being CPU bound to being memory bound, leading to a decrease in speedup. At its best, simulating typical stimulation of an Argus II over a timecourse of 500 milliseconds (at 50 microns spatial resolution and 10 ms temporal resolution) required 79 seconds of execution time. According to line profiling, most of the time is spent executing the slow convolutions (Fig. 3 D, F), thus heavily relying on the computational performance of the SciPy implementation of the Fast Fourier Transform. Due to the current design constraints (see Discussion), the software is better suited for rapid prototyping rather than real-time execution - although we aim to alleviate this in the future through GPU parallelization (via CUDA [KPL^+^12] and Dask [Das16]) and cluster computing (via Spark [Apa16]).

### Software availability and development

All code can be found at https://github.com/uwescience/pulse2percept, with up-to-date source code documentation available at https://uwescience.github.io/pulse2percept. In addition, the latest stable release is available on the Python Package Index and can be installed using pip:

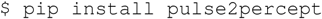

The library’s test suite can be run as follows:

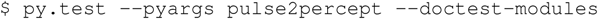

All code presented in this paper is current as of the v0.2 release.

## Discussion

We presented here an open-source, Python-based framework for modeling the visual processing in retinal prosthesis patients. This software generates a simulation of the perceptual experience of individual prosthetic users - a ‘virtual patient’.

The goal of *pulse2percept* is to provide open-source simulations that can allow any user to evaluate the perceptual experiences likely to be produced across both current and future devices. Specifically, the software is designed to meet four software design specifications:

- *Ease of use*: The intended users of this simulation include researchers and government officials who collect or assess perceptual data on prosthetic implants. These researchers are likely to be MDs rather than computer scientists, and might therefore lack technical backgrounds in computing. In the future, we plan for *pulse2percept* to become the back end of a web application similar to [KDM^+^ss].
- *Modularity*: As research continues in this field, it is likely that the underlying computational models converting electrical stimulation to patient percepts will improve. The modular design of the current release makes it easy to update individual components of the model.
- *Flexibility*: *pulse2percept* allows for rapid prototyping and integration with other analysis or visualization libraries from the Python community. It allows users to play with parameters, and use the ones that fit their desired device. Indeed, within most companies the specifications of implants currently in design is closely guarded intellectual property.
- *Extensibility*: The software can easily be extended to include custom implants, stimulation protocols, and retinal models.

As a result of these design considerations, *pulse2percept* has a number of potential uses.

Device developers can use virtual patients to get an idea of how their implant will perform even before a physical prototype has been created. This is reminiscent of the practice of virtual prototyping in other engineering fields. It becomes possible to make predictions about the perceptual consequences of individual design considerations, such as specific electrode geometries and stimulation protocols. As a result, virtual patients provide a useful tool for implant development, making it possible to rapidly predict vision across different implant configurations. We are currently collaborating with two leading manufacturers to validate the use of this software for both of these purposes.

Virtual patients can also play an important role in the wider community. To a naive viewer, simulations of prosthetic vision currently provided by manufacturers and the press might provide misleading visual outcomes, because these simulations do not take account of the substantial distortions in space and time that are observed by actual patients. On our website we provide example stimulations of real-world vision based on the *pulse2percept* virtual patient.

Prosthetic implants are expensive technology - costing roughly $100k per patient. Currently, these implants are reimbursed on a trial-by-trial basis across many countries in Europe, and are only reimbursed in a subset of states in the USA. Hence our simulations can help guide government agencies such as the FDA and Medicare in reimbursement decisions. Most importantly, these simulations can help patients, their families, and doctors make an informed choice when deciding at what stage of vision loss a prosthetic device would be helpful.

## Acknowledgments

Supported by the Washington Research Foundation Funds for Innovation in Neuroengineering and Data-Intensive Discovery (M.B.), by a grant from the Gordon & Betty Moore Foundation and the Alfred P. Sloan Foundation to the University of Washington eScience Institute Data Science Environment (M.B. and A.R.), and by the National Institutes of Health (NEI EY-012925 to G.M.B., EY-014645 to I.F.). Research credits for cloud computing were provided by Amazon Web Services.

